# G protein-coupled receptor 35 (GPR35) stimulation reduces osteoclast activity in primary human bone cells

**DOI:** 10.1101/2025.04.21.649743

**Authors:** Maria L. Price, Rachael A. Wyatt, Ana Crastin, Aqfan Jamaluddin, Rowan S. Hardy, Morten Frost, Caroline M. Gorvin

**Author notes:** **Corresponding author:** Caroline Gorvin, Department of Metabolism and Systems Science, University of Birmingham, IBR Tower, Rm 333, Birmingham, B15 2TT, United Kingdom.

## Abstract

G protein-coupled receptor 35 (GPR35) is an orphan receptor that is widely expressed in tissues including human osteoblasts and osteoclasts. Expression of the GPR35 gene and protein are downregulated in osteoporosis patients and in mouse models of the disease. *Gpr35*-knockout mice have reduced bone mass, while GPR35 agonism rescues bone loss in rodent osteoporosis models indicating that GPR35 has an important role in bone. Our previous studies demonstrated *GPR35* is expressed in human osteoclasts, and we sought to determine the receptor’s function in these cells. We differentiated human peripheral blood mononuclear cells to mature osteoclasts and assessed effects of the GPR35 synthetic agonists, TCG1001 and Zaprinast on osteoclast activity and differentiation. Both agonists stimulated significant reductions in osteoclast bone resorption and TRAP activity, and downregulated expression of *MMP9,* a gene that regulates osteoclast bone resorption. These effects were prevented by pre-incubation of cells with a GPR35-specific antagonist. To understand GPR35 signaling pathways, we measured the phosphorylation of secondary messengers known to have important roles in osteoclast activity using AlphaLISA assays. Upon GPR35 stimulation, we observed reduced phosphorylation of cSrc, which stimulates actin ring formation necessary for bone resorption, and decreased phosphorylation of Akt, CREB and NFκB that drive transcription of genes required for bone resorption. Additionally, we used chemical inhibitors and siRNA knockdown to show that GPR35 couples to Gi/o and G12/13 to stimulate these signaling pathways. Finally, we compared the ability of GPR35 agonists to suppress osteoclast activity to that of current osteoporosis drugs, denosumab and alendronic acid, and showed TRAP activity was similar suppressed under all conditions. Our findings demonstrate that GPR35 has an important inhibitory role in human osteoclast activity and have defined the signaling pathways that drive these processes. GPR35 represents a promising novel target to reduce osteoclast activity that could be exploited for osteoporosis treatments.

**Lay summary:** Expression of G protein-coupled receptor 35 (GPR35) is reduced in osteoporosis and *Gpr35*-knockout mice have reduced bone mass. Here we showed stimulation of GPR35 activates Gi/o and G12/13 signaling pathways to reduce bone resorption in human osteoclast. The anti-resorptive activity of GPR35 agonists was comparable to current osteoporosis drugs, denosumab and alendronic acid. Our findings demonstrate that GPR35 has an important inhibitory role in human osteoclast activity and have defined the signaling pathways that drive these processes. GPR35 represents a promising novel target to reduce osteoclast activity that could be exploited for osteoporosis treatments.

## Introduction

Current treatments for osteoporosis focus on prevention of bone loss or increases in bone mass using anti-resorptive, anabolic or combined anti-resorptive and anabolic drugs including bisphosphonates, parathyroid hormone-related analogues, and sclerostin inhibitors^(1)^. Although these drugs are effective in lowering fracture risk and increasing bone mass they have disadvantages including acute reactions, osteonecrosis of the jaw and atypical femur fractures for bisphosphonates^(2)^, serious cardiovascular events for the anti-sclerostin drug Romosozumab, hypercalcemia for the PTH-related analogue teriparatide^(3)^, and rapid bone loss and increased fracture risk following cessation of treatment for Denosumab^(4)^. These adverse effects indicate that additional osteoporosis therapies, which could be used alone or as part of a sequential therapy approach, are required to improve patient outcomes for osteoporosis.

G protein coupled receptors (GPCRs) are cell surface receptors that are the target of approximately one-third of all drugs approved by the US Food and Drug Administration (FDA)^(5)^, including teriparatide and abaloparatide that target the GPCR parathyroid hormone type-1 receptor (PTH1R) to treat osteoporosis. GPCRs with high expression in bone cells likely have an important role in osteoclast and/or osteoblast activity and could represent new targets for osteoporosis treatments. Our previous studies of gene expression in primary human osteoclasts using RNA-sequencing revealed that 144 GPCRs are expressed in osteoclasts^(6)^, and subsequent investigation of a subset of these GPCRs identified four receptors that reduce the activity of primary human osteoclasts^(7)^. Activation of one receptor, GPR35, significantly reduced nuclear translocation of the transcription factor nuclear factor of activated T cells-1 (NFATc1) which is essential for osteoclast differentiation and resorptive activity, and reduced bone resorption and TRAP activity in mature human osteoclasts^(7)^. However, the mechanisms by which GPR35 mediates these effects in human osteoclasts remain unknown.

GPR35 is officially designated an orphan receptor as the nature of its endogenous ligand(s) remain under investigation^(8)^. Several endogenous ligands (e.g. the tryptophan metabolite kynurenic acid^(9)^ and the serotonin metabolite 5-hydroxyindoleacetic acid (5-HIAA)^(10)^) can activate the receptor, although they have low potency or require independent verification. Therefore, studies of the physiological function of GPR35 have largely been performed with synthetic agonists such as Zaprinast or TC-G 1001^(8,11)^, while antagonists such as ML145^(12)^ have helped confirm GPR35-specificity. These studies have revealed expression of GPR35 in immune cells where it has a role in leucocyte and neutrophil recruitment to inflammatory sites^(10,13)^, the gastrointestinal tract where it may have a role in lipid metabolism and adipose tissue thermogenesis^(14)^, and in dorsal root ganglia where an anti-nociceptive role has been described^(15,16)^. GPR35 is expressed in both osteoblasts^(17)^ and osteoclasts^(18)^ and gene expression has been shown to be downregulated in humans and mice with osteoporosis^(17)^. *Gpr35^-/-^* mice have reduced bone mass due at least in part to impaired osteoblast development by reducing β-catenin activity, while activation of GPR35 in osteoporotic mice improves bone density^(17,19)^. Osteoclasts were not examined in these studies. Our recent studies suggest GPR35 stimulation impairs osteoclast activity^(7)^, however, we used Zaprinast to activate the receptor, which has been reported to inhibit phosphodiesterase (PDE)-5 and PDE6 at similar potencies to that at which it activates GPR35^(20)^. Therefore, confirmatory studies with other receptor-specific agonists, antagonists and/or gene knock-down are required to examine the mechanisms by which GPR35 regulates osteoclast activity.

Following activation, GPCRs undergo conformational changes that mediate signaling by their associated heterotrimeric Gα/β/γ proteins. Gα consists of four subfamilies: Gs that activates cAMP, Gi/o that inhibits cAMP and is pertussis toxin sensitive, Gq/11 that stimulates inositol trisphosphate (IP3) and intracellular calcium pathways and G12/13 that couples to RhoA signaling. Studies of cell-lines (e.g. HEK293) in which GPR35 is overexpressed indicate that signaling is primarily by Gi/o and G12/13^(20–23)^ and the reported CryoEM structure of GPR35 is with a chimeric G protein that has most similarity with G13^(24)^. However, it remains uncertain which Gα protein subtypes GPR35 couples to in physiologically relevant cells as this has largely been unexplored.

Here we investigated the role of GPR35 in primary human osteoclasts using two receptor agonists (TC-G 1001 and Zaprinast), a receptor antagonist (ML145) and siRNA knockdown of GPR35. We characterized the signaling pathways activated by the receptor and activity in osteoclast monocultures and osteoclast-osteoblast co-cultures. Finally, we compared how GPR35-induced reductions in osteoclast activity compares to two existing osteoclast-targeted osteoporosis drugs.

## Materials and Methods

### Compounds

Compounds were used at the following concentrations: Alendronic acid (10 µM, Cambridge Bioscience, Cambridge, UK), Denosumab (10 µM, Cambridge Bioscience, Cambridge, UK), GIP (10 nM, Bio-Techne, Abingdon, UK), GIP(3–30)NH2 (10μM, Caslo, Lyngby, Denmark), ML145 (10 µM, Tocris, Abingdon, UK), Pertussis Toxin (300 ng/mL, Tocris), TC-G 1001 (Tocris) at 10 µM for most experiments, and a concentration range in Figure 1, TUG891 (10 µM, Tocris), YM-254890 (10 µM, Cambridge Bioscience), Zaprinast (Merck, Gillingham, UK) at 10 µM for most experiments, and a concentration range in Figure 1.

**Figure 1.**
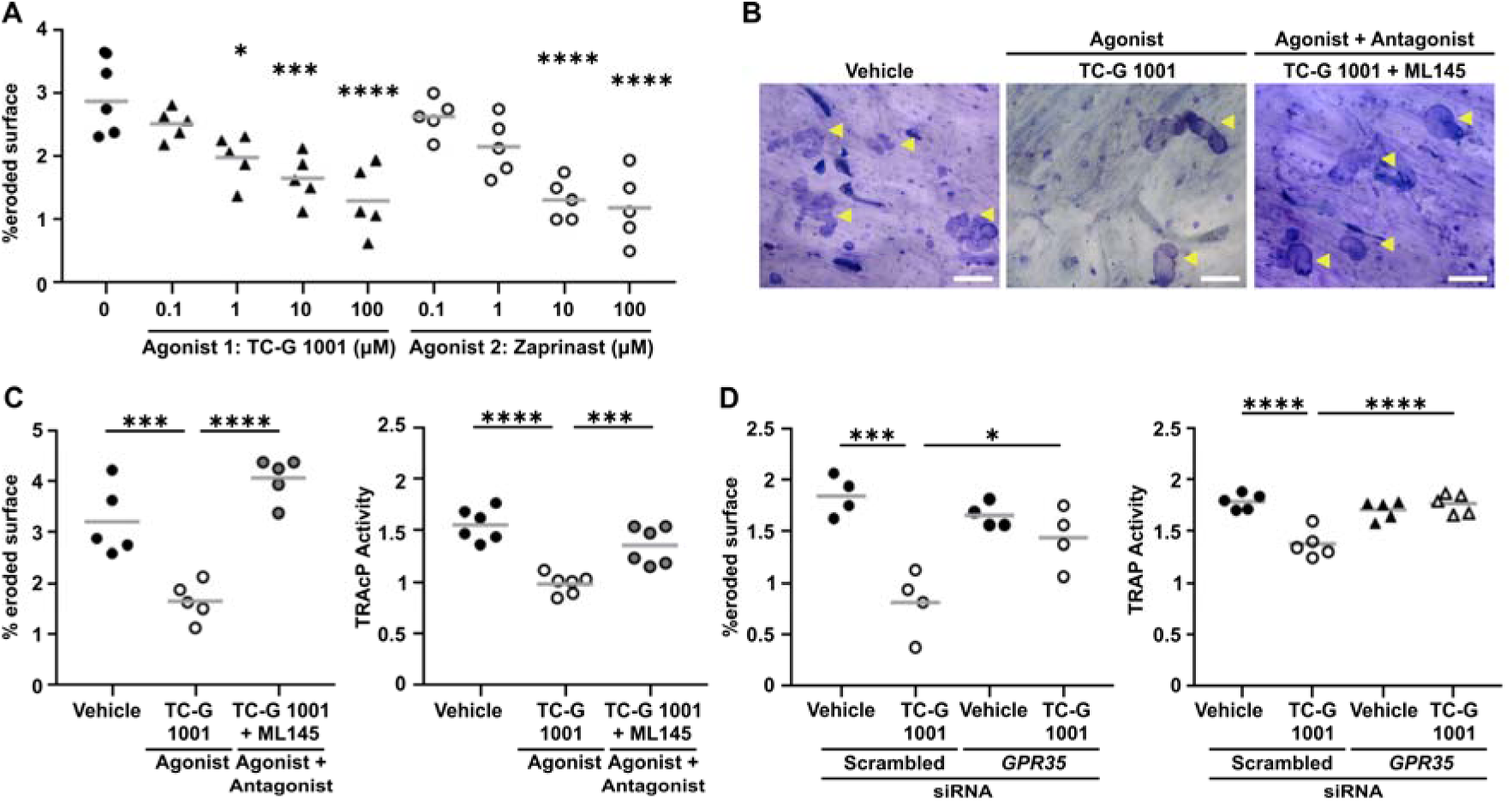
Activation of GPR35 reduces human osteoclast resorption. (**A**) Quantification of toluidine blue stained resorption pits and trenches formed on bone slices by mature osteoclasts exposed to vehicle or four concentrations of either the TC-G 1001 or Zaprinast agonist for 3 days. (**B**) Representative images of resorption pits and trenches formed on bone slices by mature osteoclasts exposed to vehicle, TC-G 1001 or TC-G 1001 and ML145 antagonist. Areas of resorption are indicated by yellow arrows. Scale bar, 100 µm. (**C**) Quantification of bone resorption and TRAP enzyme activity in conditioned media from osteoclasts treated with GPR35 agonist with and without antagonist, ML145. (**D**) Quantification of resorption pits and trenches formed on bone slices and TRAP enzyme activity by mature osteoclasts transfected with scrambled or *GPR35* siRNA and exposed to vehicle or *GPR35* agonist, TC-G 1001. Each point represents an independent donor. The gray line denotes mean in panels A, C and D. Statistical analyses were performed by one-way ANOVA with Holm-Šídák’s multiple comparisons test. ****p<0.0001, ***p<0.001, **p<0.01.

### Cell Culture

All cells were maintained at 37°C and 5% CO_2_. Primary human osteoclasts were differentiated from CD14+ monocytes, isolated from leucocyte cones obtained from anonymous blood donations from the NHS Blood and Transplant service. Approval for isolation of monocytes from human peripheral blood mononuclear cells (PBMCs) and their differentiation into osteoclasts was obtained from the local ethics committee in the UK (REC: 23/WA/0063, IRAS Project ID: 321094). Monocytes were enriched using the RosetteSep Human Monocyte Enrichment Cocktail (StemCell Technologies, Cambridge, UK) according to manufacturer’s recommendations and separated on a Ficoll-Paque gradient (VWR, Lutterworth, UK), as previously described^(25)^. Monocytes were seeded in α-minimal essential medium (αMEM, Gibco, Paisley, UK) supplemented with 10% newborn calf serum (NBCS; Gibco), 1% penicillin/ streptomycin (Fisher Scientific, Loughborough, UK) and 25 ng/mL macrophage colony-stimulating factor (M-CSF) (Bio-Techne, Abingdon, UK). Monocytes were differentiated into primary human osteoclasts over 10 days. Cells were initially stimulated with M-CSF, with media refreshed every 2-3 days. Receptor activator of nuclear factor-κβ (RANKL, 25 ng/mL, Bio-Techne) was added to cells on day 8 of differentiation to stimulate osteoclast formation. Differentiation was validated microscopically on day 10, observing the presence of multinucleated cells.

Osteoblast-like cells were differentiated from the osteoblast precursor cell-line, hMSC-TERT^(26)^, in DMEM supplemented with 10% NBCS, 1% penicillin/ streptomycin (Gibco), 10 mM β-glycerophosphate (Sigma-Aldrich, Gillingham, UK), 10 nM dexamethasone (Sigma-Aldrich), 50 µg/mL ascorbic acid (Merck), and 2mM L-glutamine (Gibco). For differentiation, media was replaced every 2-3 days and differentiation was complete on day 8, with ALP activity assays performed to confirm differentiation. hMSC-TERT were routinely tested to ensure they were mycoplasma-free using the TransDetect Luciferase Mycoplasma Detection kit (Clinisciences Ltd, Slough, UK).

### Transfection with siRNA

Trilencer-27 siRNAs were obtained from Origene Technologies (Herford, Germany) targeting three 27mer duplexes of *GNAQ* (Catalog. No. SR301847), *GNA11* (Catalog. No. SR301839), *GNA12* (Catalog. No. SR320315), *GNA13* (Catalog. No. SR307281) and *GPR35* (Catalog. No. SR301916), as well as scrambled negative controls (Catalog. No. SR30004). Primary human osteoclasts were transiently transfected with three siRNA oligos for each gene at 10 nM each, using the GenMute transfection reagent (SignaGen, Peterborough, UK). Cells were incubated with the transfection mix for 72 hours.

### Osteoclast resorption assays

Mature (day 10) osteoclasts were seeded on bovine cortical bone slices (Boneslices.com, Jelling, Denmark) in 96-well plates and allowed to settle for 1 hour. In co-culture studies, 50,000 osteoclasts per well were seeded in RANKL-free media and left to settle, before adding 12,500 osteoblast-like cells per well on top. GPCR agonists and/or antagonists were added and cells were incubated for 72 hours. Experiments were terminated by removal of media and the addition of dH_2_O to each well. Cells were removed from bone slices with a cotton swab and the bone slices were stained with toluidine blue solution (1% toluidine blue, 1% sodium borate in dH_2_O, both from Sigma-Aldrich) for 20 seconds. To visualize and quantify areas of bone resorption, blinded microscopical analyses were performed at 10x magnification on an Olympus BX53 microscope (Olympus, Tokyo, Japan), using a 10x10 counting grid (24.5 mm, Graticules Optics Ltd., Tonbridge, UK) as described^(27)^. Resorption was measured in eight fields of view spanning the bone slice and represented as the percentage eroded surface per bone slice. Sample images were taken using Olympus CellSens software (Olympus).

### Tartrate-resistant acid phosphatase 5b (TRAP) activity assays

Mature osteoclasts were seeded in 96-well plates and exposed to agonists and/or antagonists for 72 hours. Conditioned media was collected and transferred in 10 µL duplicates into a 96-well clear plate, then 90 µL TRAP solution buffer (1M acetate (Merck), 0.5% Triton X-100 (Sigma-Aldrich), 1M NaCl (Sigma-Aldrich), 10 mM EDTA (Fisher Scientific), 50 mM L-Ascorbic acid (Merck), 0.2 M disodium tartrate (Sigma-Aldrich), 82 mM 4-nitrolphenylphosphate (Sigma-Aldrich)) added to each well, prior to incubation in the dark for 30 minutes at 37°C. The reaction was stopped by adding 0.3M NaOH (Honeywell, SLS, Hessle, UK) and absorbance was measured at 405 nm on a SpectraMax ABS (Molecular Devices) or GloMax plate reader.

### AlphaLISA phosphorylation assays

Mature osteoclasts were plated in 96-well plates at a density of 50,000 cells per well in osteoclast monocultures and co-cultures. In co-culture studies, 50,000 osteoclasts per well were seeded in RANKL-free media and left to settle, before adding 12,500 osteoblast-like cells per well on top. Cells were pre-incubated with vehicle or GPR35 antagonist for 30 minutes and then exposed to vehicle or GPR35 agonist for 30 minutes. For G_q_ protein inhibition experiments, 10 µM YM-254890 was added to cells for 30 minutes with vehicle or antagonist treatments. For G_i/o_ protein inhibition experiments, cells were seeded on day 9 of differentiation and exposed to 300 ng/mL pertussis toxin for 16 hours prior to agonist and antagonist treatments. For siRNA studies, cells were transfected on day 10, then incubated for 72 hours before AlphaLISA assays were performed.

For all assays, cells were lysed in the supplied AlphaLISA buffer (PerkinElmer, Beaconsfield, UK). Lysates were transferred to white 384-well plates (OptiPlates, PerkinElmer) in duplicates and assays performed according to manufacturer’s instructions. AlphaLISA readings were made on a Pherastar FS (BMG Labtech, Aylesbury, UK) plate reader and values for phosphorylated proteins normalized to GAPDH values. The following AlphaLISA assay kits (PerkinElmer) were performed according to manufacturer’s instructions: phosphorylated forms of Akt1/2/3 (Ser473), CREB (Ser133), c-Src (Tyr419), NFkB p65 subunit (Ser536) and p38 (Thr180/Thr182) and non-phosphorylated GAPDH.

### LANCE *cAMP assays*

cAMP levels was assessed using Lance Ultra cAMP assays (Revvity, Pontyclun, UK) Mature (day 10) osteoclasts were seeded in clear 96-well plates, at a density of 50,000 cells per well, in stimulation buffer (1x Hanks Buffered Saline Solution (HBSS; Merck), 0.1% BSA (Sigma-Aldrich), 0.1% 3-isobutyl-1-methylxanthine (IBMX; Merck), 0.5 mM HEPES (Fisher Scientific)). Agonists and antagonists were diluted in stimulation buffer then added to cells and incubated for 30 minutes at 37°C. For G_i/o_ protein inhibition experiments, cells were seeded on day 9 of differentiation and exposed to PTx for 16 hours prior to agonist and antagonist treatments. Lance cAMP assays were then performed according to manufacturer’s instructions in 96-well plates. Lysates were transferred to a white 384-well plate in duplicates and HTRF readings were made at 665nm and 615 nm on a PHERAstar FS plate reader. Data was expressed as the HTRF ratio (665 nm/615 nm), values normalized to the vehicle controls.

### qPCR

To assess siRNA knockdown efficiency, mature osteoclasts were plated at 1,000,000 cells per well in a 6-well plate on day 10 of differentiation. For differentiation studies, isolated monocytes were seeded at 1,000,000 cells/well in a 35 mm dish. Osteoclasts were plated into media with compounds and media with compounds was refreshed daily, with RANKL added into the media starting on Day 8. RNA was extracted on days 3, 6 and 10 of osteoclast differentiation.

For all experiments total RNA was extracted using an RNeasy Mini Kit (Qiagen, Manchester, UK). cDNA synthesis was performed using the QuantiTect Reverse Transcription Kit (Qiagen), and qPCR performed using Quantitect primers obtained from Qiagen (Supplementary Table 1). Expression was normalized to the geometric mean of three housekeeper genes (β-actin (*ACTB*), ribosomal protein lateral stalk subunit P0 (*RPLP0*) and ubiquitin C (*UBC*)). Threshold cycle (C_T_) values were obtained from the start of the log phase on ThermoFisher Connect software and C_T_ values analyzed in Microsoft Excel using the Pfaffl method^(28)^, then data was normalized to vehicle-treated values expressed as 1.

### IP-one assays

IP-one G_q_ assays (Revvity) were used to measure inositol phosphate-1, a stable downstream metabolite of inositol phosphate 3 (IP-3). Assays were performed as described^(29)^. Mature (day 10) osteoclasts were seeded in clear 96-well plates, at 75,000 cells per well in 1x stimulation buffer (Revvity). Compounds were diluted in 1x stimulation buffer and incubated with cells for 30 minutes at 37 °C. For G_q_ protein inhibition experiments, YM-254890 was added to cells for 30 minutes with vehicle or antagonist treatments. IP-1-d2 and Anti-IP-1 Cryptate solutions (in lysis buffer) were added and incubated for 1 hour each at room temperature. Assays were performed in duplicates with 20 µL of the mix from 96-well plates transferred to white 384-well OptiPlates and the HTRF signal was read on a PHERAstar FS plate reader. Data was expressed as a ratio of 665 nm/ 620 nm then normalized to vehicle.

### Alkaline Phosphatase (ALP) Activity

Alkaline phosphatase activity was measured to confirm the presence of fully differentiated osteoblast-like cells. Fully differentiated and non-differentiated hMSC-TERT cells were seeded at 20,000 cells per well in clear 96-well plates and left to settle for 48 hours. After 48 hours, media was removed, and cells were washed with PBS then incubated with 200 µL reaction buffer (0.06 M Na_2_CO_3_ (Sigma-Aldrich), 0.04 M NaHCO_3_ (Merck), 0.1% Triton X-100 (Sigma-Aldrich), 2 mM MgSO_4_ (Fisher Scientific), 6 mM 4-NPP (Sigma-Aldrich), in water) per well for 30 min to1 hr at 37°C. To terminate the experiment, 100 µL 1M NaOH was added, then absorbance measured at 405 nm using a SpectraMax ABS (Molecular Devices) plate reader.

### Statistical Analysis

The number of experimental replicates denoted by n is indicated in figure legends. Data was exported to Microsoft Excel and statistical analyses carried out using GraphPad Prism 9. Data is presented as mean ± SEM unless otherwise stated. Normality tests (Shapiro-Wilk or D’Agostino-Pearson) were performed on all datasets to determine whether parametric or non-parametric statistical tests were appropriate. A p-value of <0.05 was considered statistically significant. Statistical analyses were performed as described in figure legends.

## Results

### Activation of GPR35 reduces human osteoclast resorption

Our previous studies have shown that activation of GPR35 with Zaprinast reduces osteoclast resorptive activity and osteoclast number^(7)^. We confirmed that the activation of GPR35 reduces osteoclast resorption in a concentration-dependent manner with two agonists, TC-G 1001 and Zaprinast (Figure 1A), and that co-treatment with the ML145 GPR35 antagonist prevented these reductions in activity (Figure 1B-C). Activation of GPR35 also reduced TRAP activity in mature osteoclasts (Figure 1C). To assess whether the TC-G 1001 agonist specifically activated GPR35, bone resorption assays and TRAP activity were repeated in the presence of a *GPR35* siRNA and compared to a scrambled siRNA. We first confirmed that the siRNA could significantly reduce *GPR35* gene expression (Supplementary Figure 1A), then assessed its effects on osteoclast activity. In the presence of scrambled siRNA, TC-G 1001 significantly reduced bone resorption and TRAP activity, which was not observed in cells transfected with *GPR35* siRNA (Figure 1D). Thus, GPR35 stimulation reduces osteoclast resorption, likely by reducing cell number.

### GPR35 activation reduces cSrc and PI3K-Akt signaling pathways in mature osteoclasts

We next sought to determine which signaling pathways GPR35 activates to mediate these anti-resorptive effects. The phosphorylation of three proteins, cSrc, Akt1/2/3 and p38, which have important roles in osteoclast activity and/or differentiation^(30–33)^ were first investigated. The phosphorylation of cSrc is important for actin ring formation in osteoclastic bone resorption^(30)^. Activation of GPR35 by both TC-G 1001 and Zaprinast reduced phosphorylation of the Tyr419 Src residue, when compared to mature osteoclasts exposed to either vehicle or antagonist (Figure 2A), consistent with reduced bone resorption by GPR35. The phosphorylation of Akt can promote osteoclast resorption or osteoclast differentiation^(31,34)^. Phosphorylation of Akt1/2/3 (pAkt) was significantly reduced in mature osteoclasts exposed to GPR35 agonists, with effects abolished in cells pre-treated with ML145 (Figure 2B). Osteoclast differentiation can also be induced by the p38 mitogen-activated protein kinase (MAPK) pathway^(32,33)^. However, GPR35 activation did not affect phosphorylation of p38 (Figure 2C). Pre-treatment of osteoclasts with the *GPR35* siRNA prevented the reduction in phosphorylated cSrc and Akt1/2/3, further demonstrating that these responses are GPR35 mediated (Figure 2D-E).

**Figure 2.**
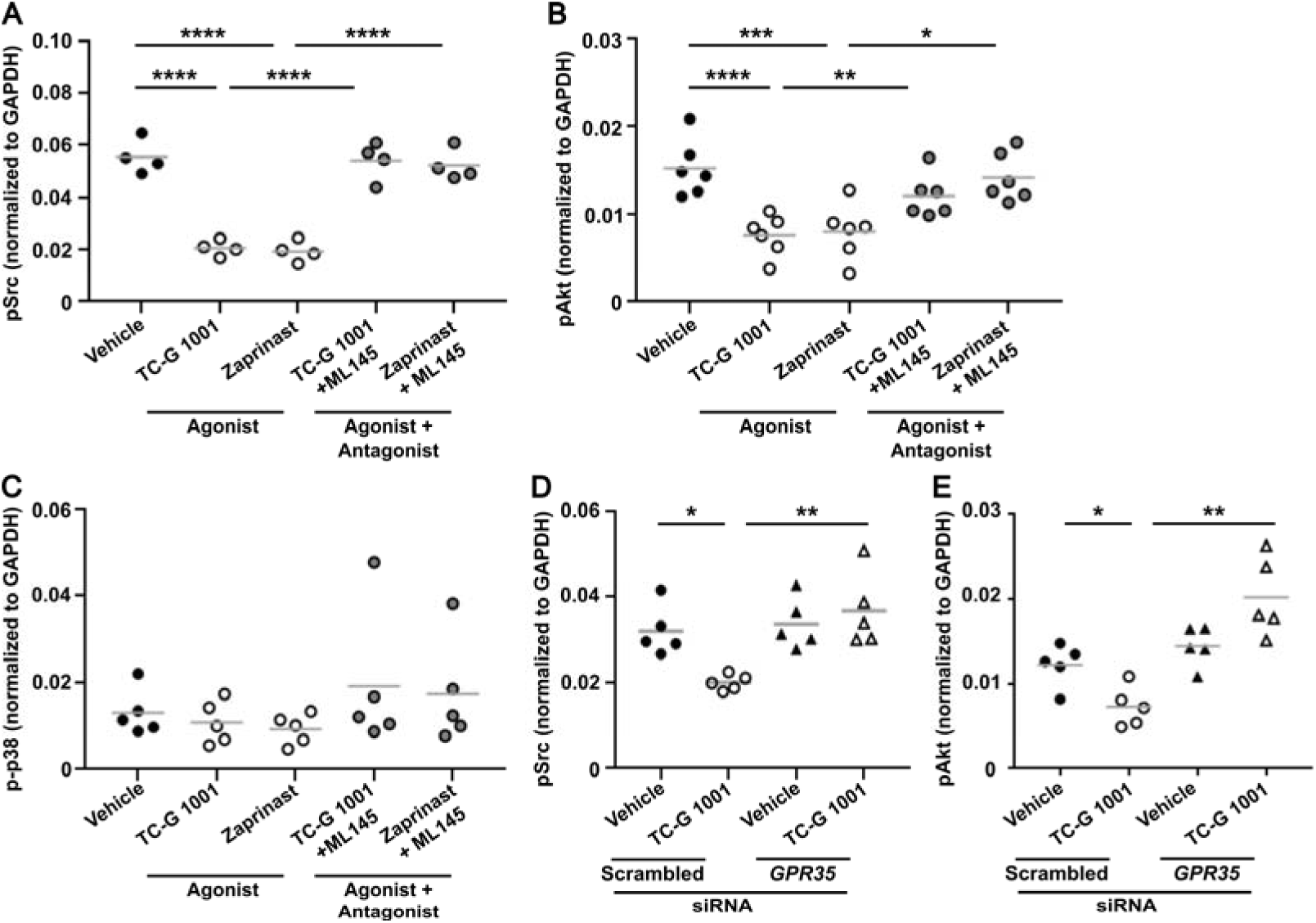
GPR35 activation reduces cSrc and PI3K-Akt signaling pathways in mature osteoclasts. Quantification of phosphorylated (**A**) cSrc (pSrc), (**B**) Akt1/2/3 (pAkt), and (**C**) p38 (p-p38) measured by AlphaLISA in osteoclasts exposed to vehicle or GPR35 agonists with or without GPR35 antagonist ML145. (**D-E**) Quantification of (D) cSrc and (E) Akt1/2/3 phosphorylation in mature osteoclasts transfected with scrambled or *GPR35* siRNA exposed to vehicle or *GPR35* agonist, TC-G 1001. Data in all panels was normalized to GAPDH as a housekeeper control. Each point represents an independent donor in all panels. The gray line denotes mean in panels A, B, D and E and median in panel C. Statistical analyses were performed by one-way ANOVA with Holm-Šídák’s multiple comparisons test in panels A, B, D and E and Kruskal-Wallis with Dunn’s multiple comparisons test in C. ****p<0.0001, ***p<0.001, **p<0.01, *p<0.05.

### GPR35 activation reduces transcription factor activation and osteoclast-specific gene expression

A number of signaling pathways are known to be activated downstream of c-Src and PI3K-Akt including the nuclear factor κB (NFκB)^(35–37)^ and cAMP response element binding protein (CREB)^(38)^. Phosphorylation of the p65 NFκB subunit and CREB protein were reduced by GPR35 activation with TC-G 1001 and Zaprinast when compared to mature osteoclasts exposed to vehicle or antagonist with agonist (Figure 3A-B). NFκB and CREB activate NFATc1 nuclear translocation which induces the expression of genes with roles in osteoclast differentiation and activation. As GPR35 activation suppresses NFκB and CREB phosphorylation and we have previously shown GPR35 reduces NFATc1 nuclear translocation^(7)^, we assessed the expression of two known NFATc1 target genes, cathepsin K (*CTSK*) and matrix metallopeptidase 9 (*MMP9*). To assess the effect of GPR35 stimulation on differentiation, cells were exposed to GPR35 agonists from monocyte isolation on day 1 and gene expression assessed on days 3, 6 and 10 of osteoclast differentiation. *MMP9* expression was significantly reduced on days 6 and 10 by GPR35 activation when compared to cells exposed to vehicle (Figure 3C). No difference in *CTSK* gene expression was detected (Figure 3D). Thus, GPR35 activation may impact matrix organization rather than direct effects on resorption.

**Figure 3.**
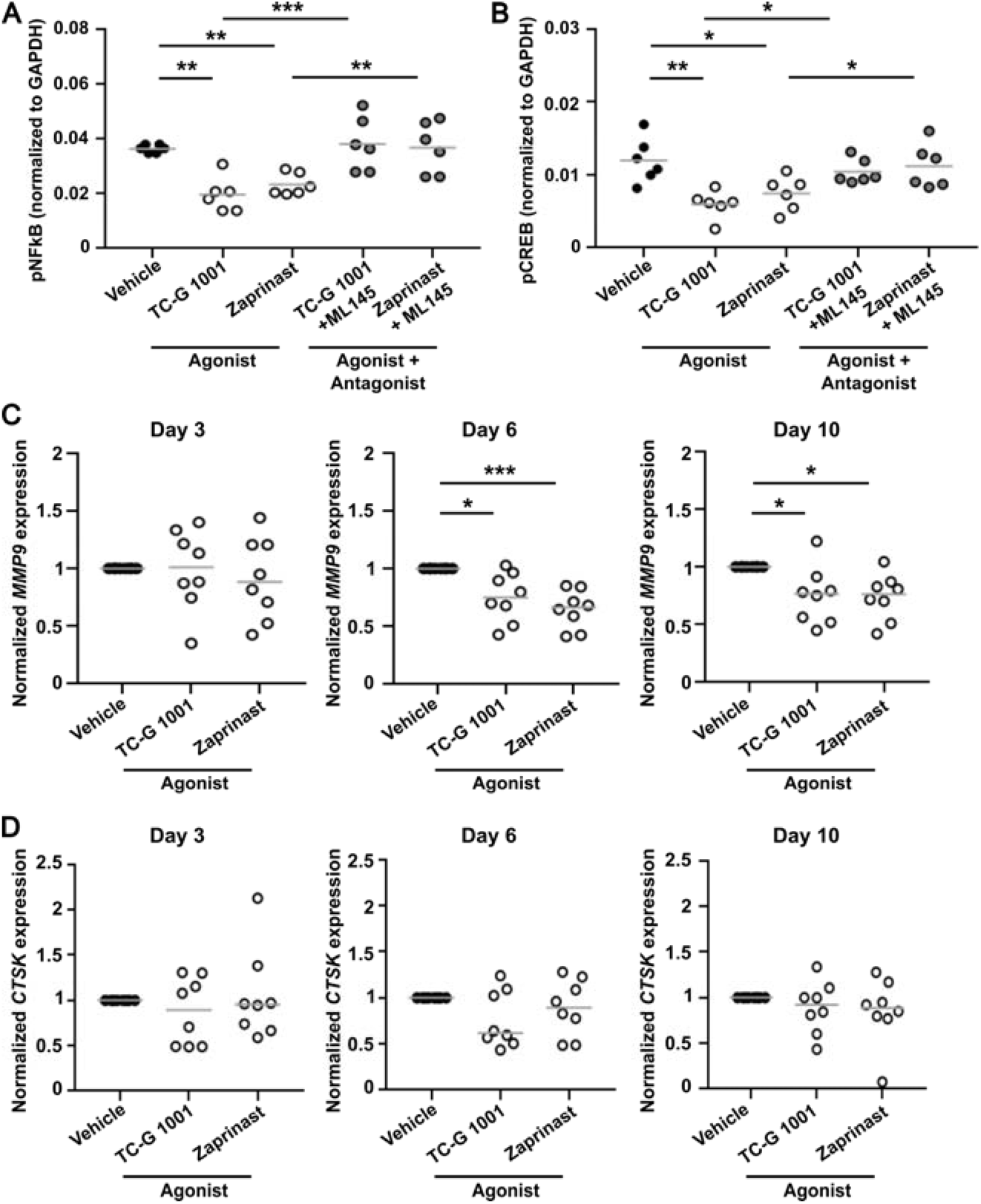
GPR35 activation reduces NFκB and CREB signaling and suppresses *MMP9* expression. Quantification of (**A**) phosphorylated NFκB (pNFκB) or (**B**) phosphorylated CREB (pCREB) in osteoclasts exposed to vehicle or GPR35 agonists with or without GPR35 antagonist ML145. Gene expression of (**C**) *MMP9* and (**D**) *CTSK* in primary human osteoclasts following treatment with GPR35 agonists, TC-G 1001 and Zaprinast on day 3, day 6 and day 10 of osteoclast differentiation determined by qPCR. Each point represents an independent donor. The gray line denotes mean in A-B and median in C-D. Statistical analyses were performed by one-way ANOVA with Holm-Šídák’s multiple comparisons test in A-B and by Kruskal-Wallis test with Dunn’s multiple comparisons test in C-D. ****p<0.0001, ***p<0.001, **p<0.01, *p<0.05.

### Effects of GPR35 on bone are maintained in osteoclast-osteoblast co-cultures

In osteoclast monocultures, cells are supplemented with RANKL to induce osteoclast differentiation. To determine whether GPR35-induced reductions in osteoclast resorption are maintained in an environment in which osteoclasts respond to osteoblast-mediated RANKL secretion, the role of GPR35 was investigated in osteoclasts grown with osteoblasts differentiated from hMSC-TERT cells^(26)^. The differentiation of hMSC-TERT cells to osteoblast-like cells was confirmed by measuring alkaline phosphatase activity (Supplementary Figure 2). Mature osteoclasts were cultured with differentiated hMSC-TERT osteoblast-like cells to examine GPR35 activity in osteoclast-osteoblast co-cultures. Exposure of co-cultures to the GPR35 agonists TC-G 1001 and Zaprinast for 72 hours impaired bone resorption and TRAP activity in co-cultures when compared to cells exposed to vehicle or agonist and antagonist (Figure 4A-B). GPR35 agonists also reduced phosphorylated Src, Akt1/2/3, NFkB and CREB concentrations in osteoclast-osteoblast co-cultures, while cells with antagonist were not significantly different to cells exposed to vehicle (Figure 4C-F). Thus, GPR35-mediated effects on osteoclast activity are retained in osteoclast-osteoblast co-cultures.

**Figure 4.**
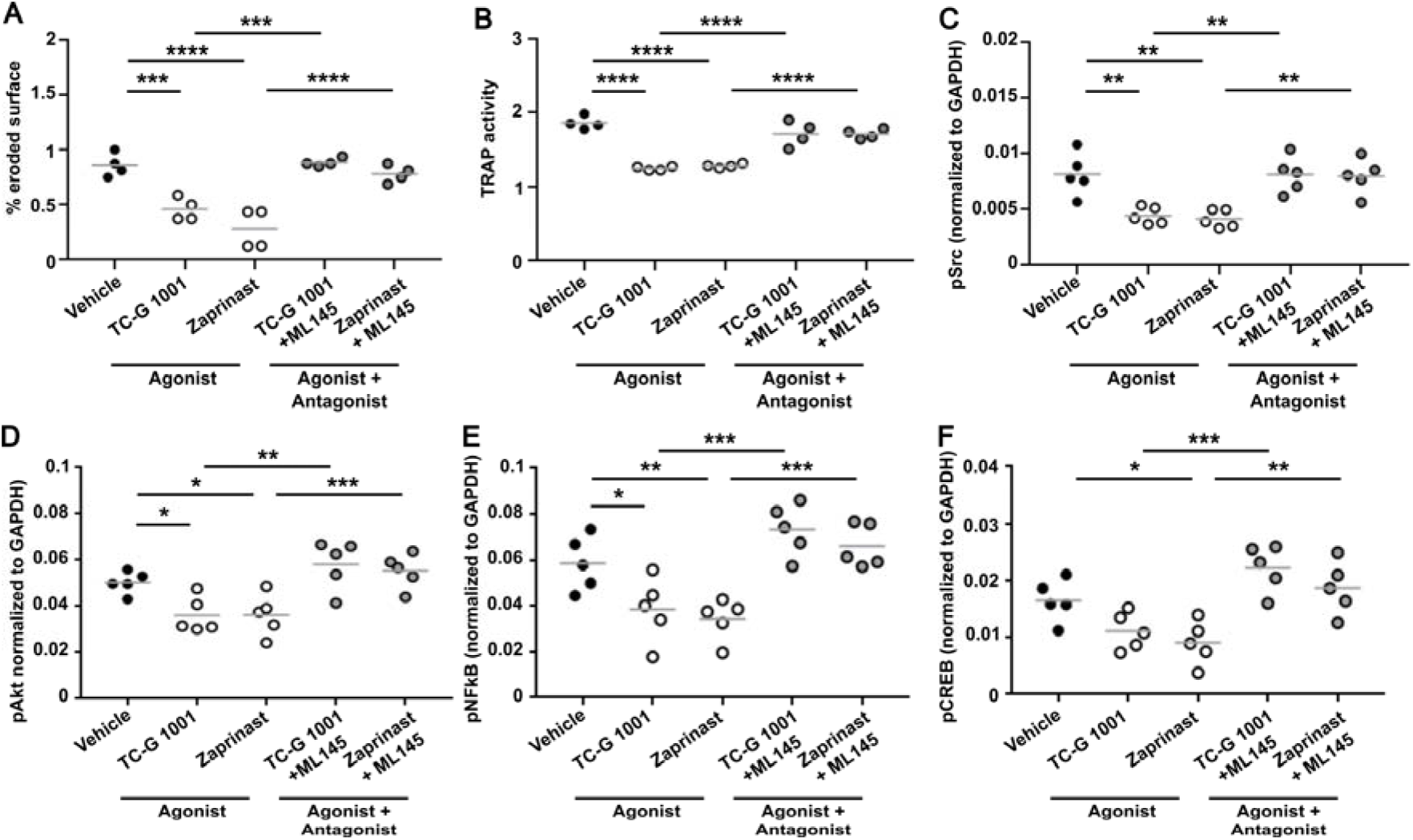
GPR35 mediated effects on bone resorption and signaling are maintained in osteoclast-osteoblast co-cultures. (**A**) Quantification of toluidine blue stained resorption pits and trenches formed on bone slices by osteoclast-osteoblast co-cultures exposed to vehicle, TC-G 1001 or TC-G 1001 and the ML145 antagonist (**B**) Quantification of TRAP enzyme activity in conditioned media from osteoclast-osteoblast co-cultures exposed to GPR35 agonists or agonists with antagonist. (**C**-**F**) Quantification of (C) phosphorylated Src (pSrc), (D) phosphorylated Akt1/2/3 (pAkt), (E) phosphorylated NFκB, (F) phosphorylated CREB in osteoclast-osteoblast co-cultures exposed to GPR35 agonists or agonists with antagonist, measured by AlphaLISA. Each point represents an independent donor. The gray line denotes mean. Statistical analyses were performed by one-way ANOVA with Holm-Šídák’s multiple comparisons test for A-B and with Dunnett’s multiple comparisons test for C-F. ****p<0.0001, ***p<0.001, **p<0.01, *p<0.05.

### GPR35 couples to G_i/o_ and G_12/13_ proteins in primary human osteoclasts

Previous studies in HEK293 and U20S cells showed overexpression of GPR35 increases signaling by G12/13 and Gi/o^(15,20–22,39)^. To determine whether endogenous GPR35 functional effects in primary human osteoclasts involve these same signaling pathways we repeated the pSrc assays in the presence of inhibitors and/or siRNAs targeting each G protein family. Signaling by cAMP was first assessed as Gs and Gi/o converge on this pathway. Mature osteoclasts were exposed to TC-G 1001 and Zaprinast and cAMP accumulation assessed by LANCE assays. There was no significant change in cAMP concentrations upon stimulation of GPR35 (Figure 5A), while increases in cAMP were detected in cells exposed to GIP, which is known to activate Gs-mediated signaling^(27)^ (Supplementary Figure 3). Therefore, GPR35 is unlikely to signal by Gs pathways. To assess Gi/o signaling, mature osteoclasts were exposed to forskolin to elevate cAMP concentrations, then responses to GPR35 agonists and antagonists assessed. TC-G 1001 and Zaprinast significantly reduced forskolin-induced cAMP responses when compared to cells exposed to vehicle or antagonist, indicating that GPR35 signals by Gi/o pathways in primary human osteoclasts (Figure 5B). To determine whether GPR35 coupling to Gi/o contributes to signaling pathways that are known to be important for bone resorption, pSrc assays were performed in the presence of the Gi/o antagonist pertussis toxin (PTx). Pre-treatment of cells with PTx abolished GPR35-mediated effects on pSrc (Figure 5C).

**Figure 5.**
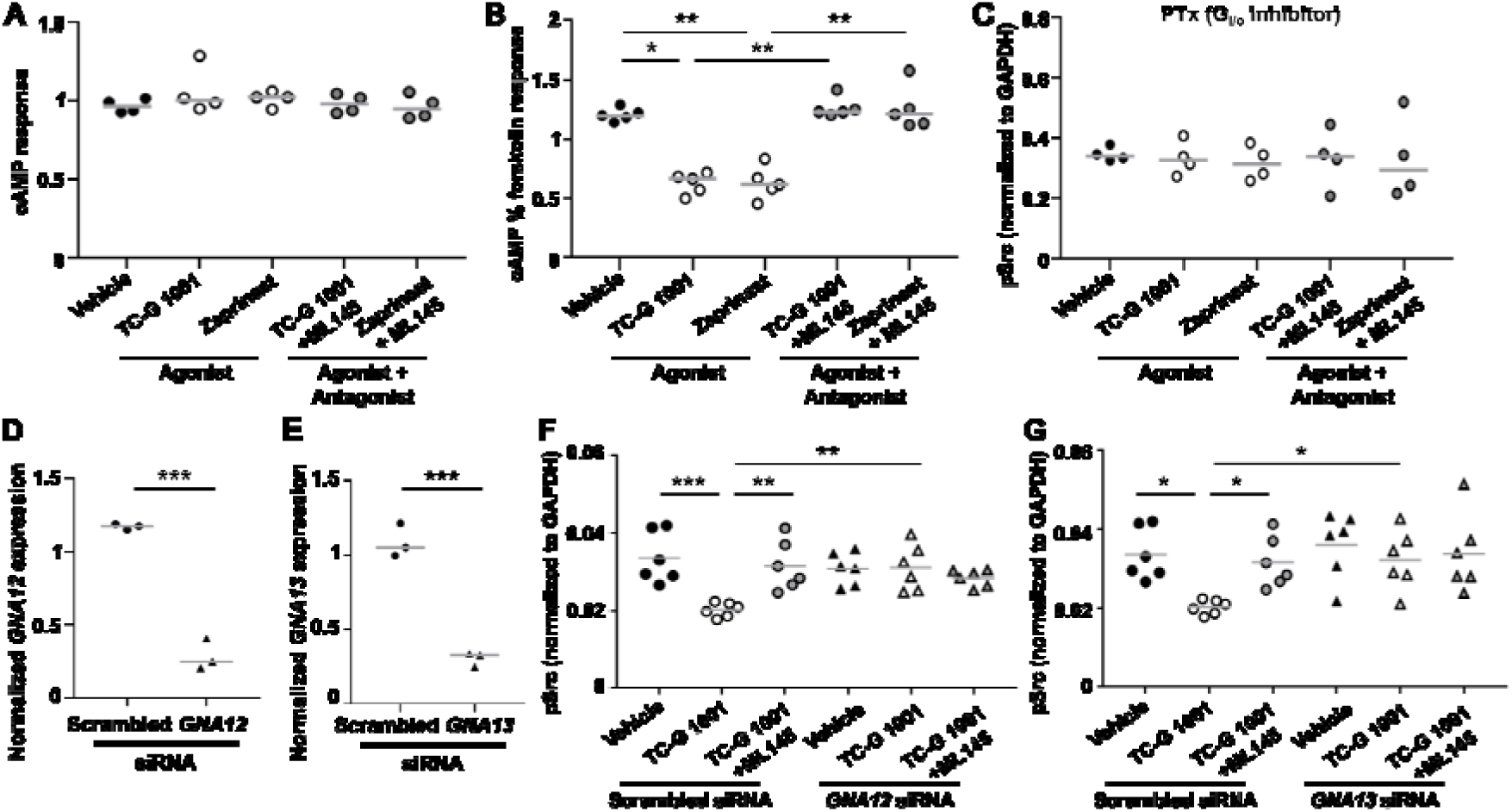
GPR35 couples to G_i/o_ and G_12/13_ proteins in primary human osteoclasts. (**A**) Quantification of cAMP levels by LANCE assays to test Gs signaling in primary human osteoclasts upon exposure to vehicle or GPR35 agonists with or without GPR35 antagonist ML145. (**B**) Suppression of forskolin-induced cAMP concentrations to test Gi/o signaling in osteoclasts exposed to vehicle, GPR35 agonists or agonists with antagonist. cAMP levels were normalized to average response by donor. (**C**) Quantification of GPR35-mediated pSrc concentrations in osteoclasts exposed to the Gi/o inhibitor pertussis toxin (PTx). (D-E) Gene expression of (D) *GNA12* and (E) *GNA13* in mature osteoclasts exposed to scrambled or targeted siRNAs determined by qPCR. Data was normalized to the geometric mean of three housekeeper genes (*ACTB, RPLP0, UBC*). (**F**-**G**) Quantification of GPR35-mediated pSrc concentrations in osteoclasts exposed to (F) *GNA12* siRNA and (G) *GNA13* siRNA. The gray line denotes median in panels A-B and mean in panels C-G. Statistical analysis by Kruskal-Wallis test with Dunn’s multiple comparisons test in panels A-B, one-way ANOVA with Holm-Šídák’s or Dunnett’s multiple comparisons test for panel C, F, G, unpaired t-test in D-E. ***p<0.001, **p<0.01, *p<0.05.

To investigate G12/13 pathways, osteoclasts were transfected with siRNAs targeting either *GNA12, GNA13* or a control siRNA. The ability of these siRNAs to reduce *GNA12* and *GNA13* gene expression was demonstrated in mature human osteoclasts by qPCR (Figure 5D-E). Assays to assess pSrc signaling with GPR35 agonists were then performed in the presence of each siRNA. Knockdown of *GNA12* and *GNA13* abolished the effects of TC-G 1001 on pSrc in mature osteoclasts (Figure 5F-G). Therefore, G12 and G13 may contribute to pSrc signaling in response to GPR35 stimulation.

The Gq/11 pathway was initially examined by measuring IP-1 concentrations in mature osteoclasts exposed to TC-G 1001 and Zaprinast. No significant differences were identified when compared to vehicle treated cells (Figure 6A), although the FFAR4 agonist, TUG891, which we have previously shown to activate Gq/11-mediated responses^(18)^, did increase IP-1 concentrations (Supplementary Figure 3). To verify these findings we assessed GPR35-mediated effects on pSrc signaling in mature osteoclasts exposed to the Gq/11 inhibitor YM-254890 and siRNAs targeting *GNAQ* and *GNA11*. TC-G 1001 and Zaprinast retained their ability to reduce pSrc signaling in the presence of YM-254890 and the *GNAQ* and *GNA11* siRNAs indicating that it is unlikely that GPR35 couples to Gq/11 in mature osteoclasts (Figure 6B-D).

**Figure 6.**
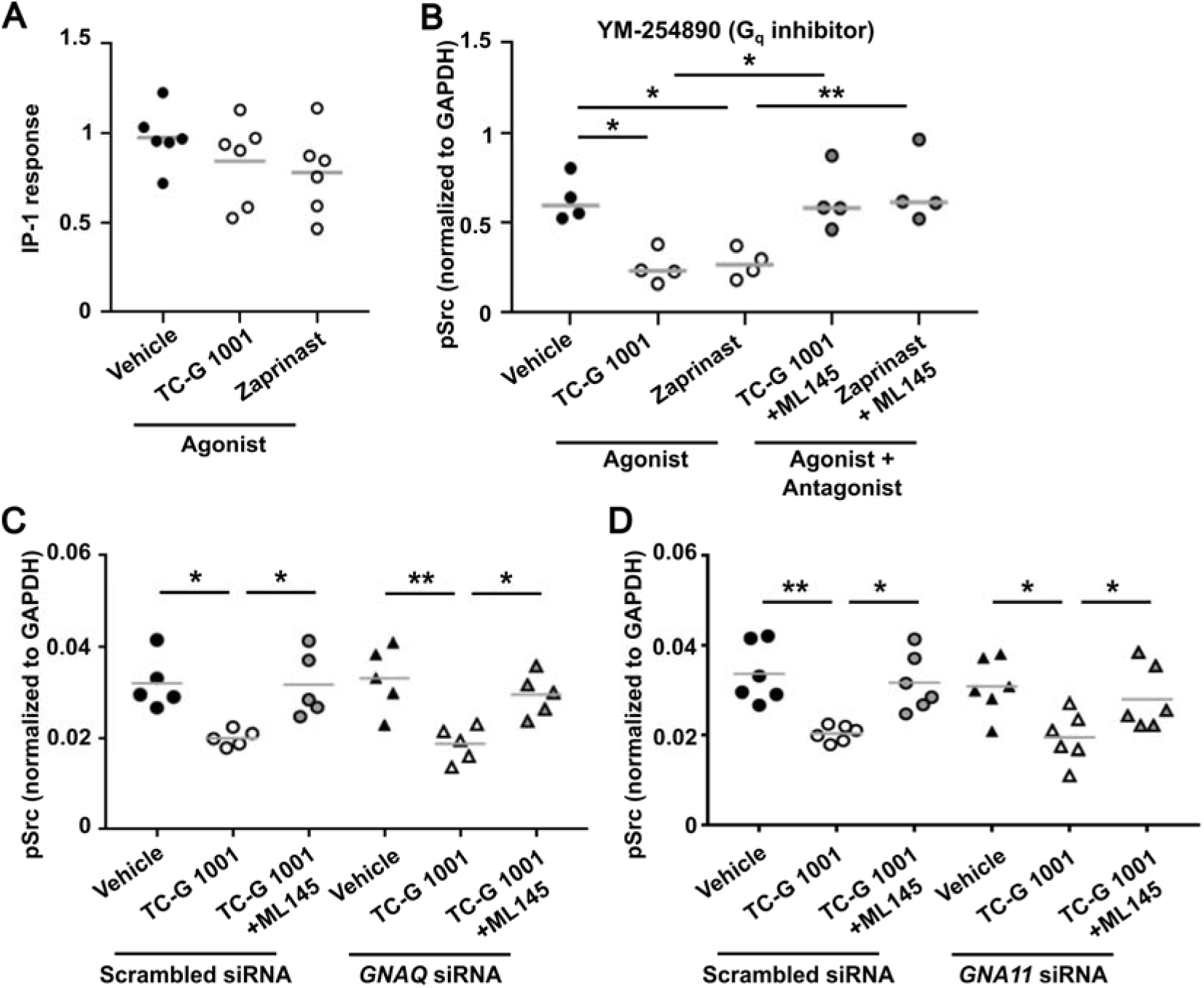
GPR35 does not couple to G_q/11_ proteins in primary human osteoclasts. (**A**) IP-1 concentrations measured by IP-one assays in primary human osteoclasts exposed to vehicle or the GPR35 agonists, TC-G 1001 and Zaprinast. (**B-D**) Quantification of GPR35-mediated pSrc concentrations in osteoclasts exposed to (B) the Gq/11 inhibitor YM-254890, (C) *GNAQ* siRNA and (D) *GNA11* siRNA and treated to agonist TC-G 1001 with or without GPR35 antagonist ML145. Each point represents an independent donor. The gray line denotes mean. Statistical analysis by one-way ANOVA with Holm-Šídák’s or Dunnett’s multiple comparisons test for panel B-D. ****p<0.0001, ***p<0.001, **p<0.01, *p<0.05.

### GPR35 agonists perform as well as current osteoporosis drugs *in vitro*

Previous studies have indicated that GPR35 expression is reduced in bone marrow mesenchymal stem cells of humans and mice with osteoporosis^(17)^, which could limit the feasibility of targeting GPR35. However, our previous studies have shown GPR35 expression is high in monocytes^(18)^ and we have shown osteoclasts can respond to GPR35 agonists (Figure 1-6). To determine whether GPR35 stimulation affects *GPR35* expression we exposed cells to Zaprinast or TC-G 1001 for 6 and 10 days of osteoclast differentiation and compared expression by qPCR relative to cells exposed to vehicle. This showed a significant increase in *GPR35* expression at both time-points (Figure 7A).

**Figure 7.**
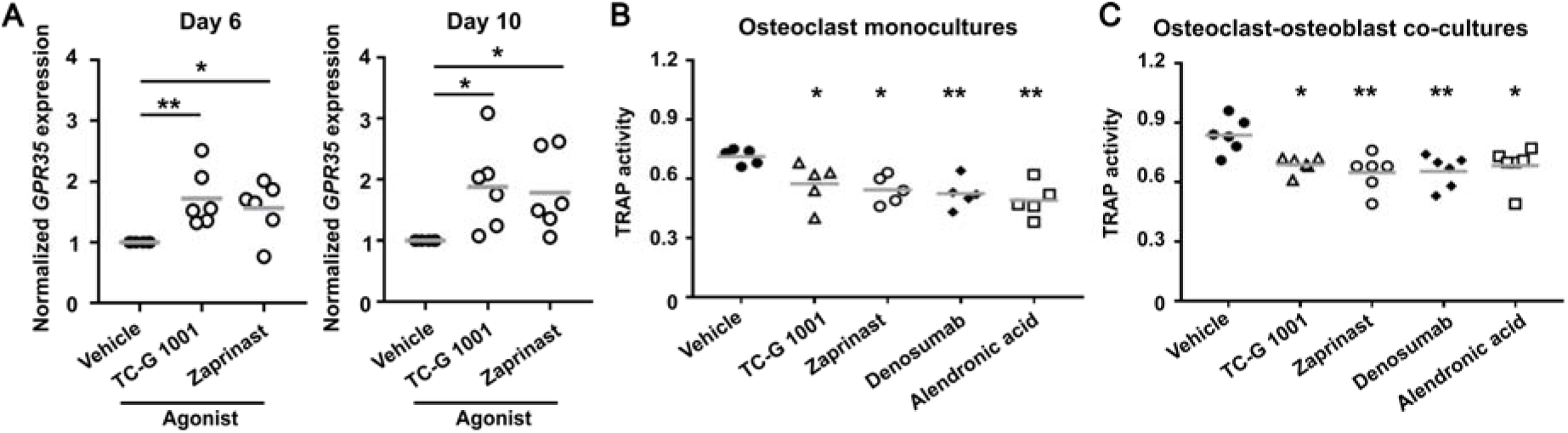
Stimulation of GPR35 reduces osteoclast activity similarly to current osteoporosis drugs. (**A**) Gene expression of *GPR35* in primary human osteoclasts following treatment with GPR35 agonists, TC-G 1001 and Zaprinast on day 6 and day 10 of osteoclast differentiation determined by qPCR. Data was normalized to the geometric mean of three housekeeper genes (*ACTB, RPLP0, UBC*). (**B**-**C**) Quantification of TRAP in conditioned media from (B) osteoclast monocultures or (C) osteoclast-osteoblast co-cultures treated with GPR35 agonists, 10 µM denosumab or 10 µM alendronic acid. Each point represents an independent donor in all panels. The gray line denotes mean. Statistical analyses were performed by one-way ANOVA with Holm-Šídák’s multiple comparisons test with comparisons to vehicle shown in B and C. **p<0.01, *p<0.05.

Finally, we compared the ability of the GPR35 agonists to reduce osteoclast activity to two currently available osteoporosis drugs, denosumab and alendronic acid (alendronate). TRAP activity assays were performed in osteoclast monocultures and osteoclast-osteoblast co-cultures, with cells exposed to all compounds for 72 hours. GPR35 agonists reduced TRAP activity to similar levels to denosumab and alendronic acid (Figure 7B-C).

## Discussion

We have demonstrated that stimulation of GPR35 in primary human osteoclasts impairs osteoclast activity and may be a viable target in osteoporosis therapies. We provide several lines of evidence that direct stimulation of GPR35 is responsible for these effects. Firstly, we used two synthetic agonists, TC-G 1001 and Zaprinast, that have been reported to activate GPR35 in multiple studies. While previous studies indicate that Zaprinast has off-target effects on phosphodiesterase-5 and -6^(40)^, PDE5 is not expressed and PDE6 is expressed at very low concentrations in mature osteoclasts^(6)^, and use of a GPR35-specific antagonist in all studies increases confidence that our findings are mediated by GPR35 activity. Moreover, Zaprinast and TC-G 1001-mediated effects were abolished in cells depleted of *GPR35* by siRNA. Furthermore, our findings are consistent with animal studies that show kynurenic acid, a putative ligand for GPR35, reduces TRAP activity in femurs of ovariectomized mice^(41)^.

Our studies provide a comprehensive assessment of the signaling pathways activated by GPR35 within physiologically-relevant cells and insights into the mechanisms by which receptor stimulation impairs osteoclast activity. Thus, GPR35 impairs the phosphorylation of Src, which facilitates actin ring formation in osteoclastic bone resorption^(30)^, reduces phosphorylation of Akt1/2/3, which promotes osteoclast resorption and osteoclast differentiation^(31,34)^, and subsequently reduces activation and nuclear translocation of downstream transcription factors including CREB, NFκB and NFATc1^(7)^ that regulate the expression of genes that promote osteoclast differentiation and function^(31,35–38)^. Consistent with these signaling pathways we have previously shown reductions in TRAP activity and cell viability^(7)^, and here show reduced bone resorption and expression of *MMP9*, which has an important role in osteoclast differentiation and function downstream of NFATc1^(42)^. In contrast, we did not observe a change in *CTSK* expression, although kynurenic acid has been shown to reduce *CTSK* in ovariectomized mice^(41)^. This may be because there is greater variation in human samples or that estrogen deficiency in ovariectomized mice influences gene expression. Our ethics approval does not allow us to determine the age or gender of the individuals that donate blood samples and therefore we do not know whether sex hormones may have influenced our findings.

These studies demonstrate that GPR35 couples predominantly to G12/13 and Gi/o signaling pathways following receptor activation consistent with findings in transfected non-osteoclast cell-lines^(20–22,43)^ and the reported CryoEM structure of GPR35 bound to a chimeric G13 protein^(24)^. In contrast, osteoclast-GPR35 does not appear to couple to Gq/11 pathways that have been reported in one study of transfected HEK293 cells^(11)^. As we did not examine constitutive activation of the receptor we do not know whether GPR35 is capable of signaling by other pathways in osteoclasts in the ligand-free state. However, other studies suggest constitutive activity is largely mediated by G12/13^(23,43)^. Few studies have examined the G protein selectivity of endogenous GPR35. Studies of GPR35-dependent chemotactic responses in B lymphoma cells and neutrophils have shown these functions are pertussis toxin sensitive^(10)^ and treatment of HepG2 cells (a model of hepatocytes) with the Rho-kinase inhibitor Y27632 abolished GPR35-mediated signaling^(40)^. It is likely that GPR35 has the capacity to signal by both G12/13 and Gi/o, and that the tissue specific context (e.g. G protein or effector expression levels) determines which pathway is preferentially activated.

GPR35 agonists reduced TRAP activity in osteoclasts and co-cultures to similar levels to denosumab and alendronic acid which are currently used to treat osteoporosis. Previous studies in mice showed that GPR35 activation improves bone density in osteoporosis by direct actions on osteoblasts, but neither of these studies assessed osteoclast effects^(17,19)^. Therefore, GPR35 agonism could have both anti-resorptive and anabolic effects on bone and is a promising candidate for osteoporosis treatment. Although previous studies indicate that expression of the *GPR35* gene is reduced in humans and mice with osteoporosis, stimulation of monocytes with GPR35 agonists significantly increased *GPR35* expression in pre-osteoclasts and mature osteoclasts indicating that the receptor is a viable target. Detailed studies of GPR35 agonists in comparison to current osteoporosis drugs in mice could provide further evidence for these findings, although such studies would likely need to be performed in mice with humanized *GPR35* as most GPR35 agonists have lower potency at mouse GPR35 when compared to human^(8,40)^.

GPR35 is widely expressed in immune cells, cardiomyocytes, hepatocytes, the gastrointestinal tract and dorsal root ganglia^(10,13–16,20,44)^, and off-target effects on these cells must be considered if GPR35 is to be targeted in osteoporosis. GPR35 agonism generally has beneficial effects on these tissues including protective roles against cardiac ischemia, anti-nociception, anti-inflammatory roles in models of colitis and prevention and reversal of lipid accumulation^(10,13–16,20,44)^. It seems unlikely that these will have detrimental adverse effects, and many will be beneficial in the generally older population with osteoporosis who may be living with multiple comorbidities. However, immune responses may need to be monitored in these patients. As GPR35 agonism is increasingly investigated to treat inflammatory bowel disease, non-alcoholic fatty liver disease and cardiac ischemia^(8,40,44)^ it is likely that improved highly-potent small molecules will be designed to target the receptor and these compounds could have efficacy in treating osteoporosis.

In summary we have demonstrated that GPR35 has an anti-resorptive role in human osteoclasts by reducing multiple phosphorylated proteins downstream of G12/13 and Gi/o signaling pathways. Drugs that stimulate GPR35 could have anti-resorptive and anabolic effects on bone and represent a viable new target for osteoporosis treatments.

## Author contributions

Methodology: MLP, CMG

Investigation: MLP, RAW, AJ, AC, RSH, CMG

Supervision: MF, CMG

Writing – original draft: CMG

Writing – review and editing: All authors

## Data and materials availability

All data needed to evaluate the conclusions in the paper are present in the paper and/or the Supplementary Materials. Enhanced expression of *GPR35* in mature osteoclasts was first identified in RNA-sequencing data generated from cultured primary osteoclasts derived from eight human donors have been deposited in the Gene Expression Omnibus (GEO) database under accession code GSE246769. Processed scRNA-seq data are available at the open science framework: https://osf.io/9xys4/.

## Funding

This work was supported by: the Novo Nordisk Foundation (Grant number NNF18OC0055047 to MF), a Sir Henry Dale Fellowship jointly funded by the Wellcome Trust and the Royal Society (Grant Number 224155/Z/21/Z to CMG), and The National Institute for Health and Care Research (NIHR) Biomedical Research Centre (BRC) at the University of Birmingham (funding for AC).

## Declaration of Interest

The authors have no conflicts to disclose.

## Ethics approval

Ethics for differentiation of human osteoclasts from blood samples: Research Ethics Committee (REC): Wales REC 7, REC: 23/WA/0063, IRAS Project ID: 321094.

## Notes

### Competing Interest Statement

The authors have declared no competing interest.

## References

1. Park Y, Sato T, Lee J. Functional and analytical recapitulation of osteoclast biology on demineralized bone paper. Nat Commun. Dec 7 2023;14(1):8092.

2. Khosla S, Bilezikian JP, Dempster DW, Lewiecki EM, Miller PD, Neer RM, et al. Benefits and risks of bisphosphonate therapy for osteoporosis. J Clin Endocrinol Metab. Jul 2012;97(7):2272–82.

3. Wen MT, Li JC, Lu BW, Shao HR, Ling PX, Liu F, et al. Indications and adverse events of teriparatide: based on FDA adverse event reporting system (FAERS). Front Pharmacol. 2024;15:1391356.

4. Ferrari S, Langdahl B. Mechanisms underlying the long-term and withdrawal effects of denosumab therapy on bone. Nat Rev Rheumatol. May 2023;19(5):307–17.

5. Hauser AS, Attwood MM, Rask-Andersen M, Schioth HB, Gloriam DE. Trends in GPCR drug discovery: new agents, targets and indications. Nat Rev Drug Discov. Dec 2017;16(12):829–42.

6. Hansen MS, Madsen K, Price M, Søe K, Omata Y, Zaiss MM, et al. Transcriptional reprogramming during human osteoclast differentiation identifies regulators of osteoclast activity. Bone Research 2024 12:1. 2024;12(1):1–19.

7. Price ML, Wyatt RA, Correia J, Areej Z, Hinds M, Crastin A, et al. Identification of anti-resorptive GPCRs by high-content imaging in human osteoclasts. J Mol Endocrinol. Apr 1 2025.

8. Milligan G. GPR35: from enigma to therapeutic target. Trends Pharmacol Sci. May 2023;44(5):263–73.

9. Wang J, Simonavicius N, Wu X, Swaminath G, Reagan J, Tian H, et al. Kynurenic acid as a ligand for orphan G protein-coupled receptor GPR35. J Biol Chem. Aug 4 2006;281(31):22021–8.

10. De Giovanni M, Tam H, Valet C, Xu Y, Looney MR, Cyster JG. GPR35 promotes neutrophil recruitment in response to serotonin metabolite 5-HIAA. Cell. Mar 3 2022;185(5):815–30 e19.

11. Neetoo-Isseljee Z, MacKenzie AE, Southern C, Jerman J, McIver EG, Harries N, et al. High-throughput identification and characterization of novel, species-selective GPR35 agonists. J Pharmacol Exp Ther. Mar 2013;344(3):568–78.

12. Heynen-Genel S, Dahl R, Shi S, Sauer M, Hariharan S, Sergienko E, et al. Selective GPR35 Antagonists - Probe 3. Probe Reports from the NIH Molecular Libraries Program. Bethesda (MD)2010.

13. Barth MC, Ahluwalia N, Anderson TJ, Hardy GJ, Sinha S, Alvarez-Cardona JA, et al. Kynurenic acid triggers firm arrest of leukocytes to vascular endothelium under flow conditions. J Biol Chem. Jul 17 2009;284(29):19189–95.

14. Agudelo LZ, Ferreira DMS, Cervenka I, Bryzgalova G, Dadvar S, Jannig PR, et al. Kynurenic Acid and Gpr35 Regulate Adipose Tissue Energy Homeostasis and Inflammation. Cell Metab. Feb 6 2018;27(2):378–92 e5.

15. Zhao P, Sharir H, Kapur A, Cowan A, Geller EB, Adler MW, et al. Targeting of the orphan receptor GPR35 by pamoic acid: a potent activator of extracellular signal-regulated kinase and beta-arrestin2 with antinociceptive activity. Mol Pharmacol. Oct 2010;78(4):560–8.

16. Cosi C, Mannaioni G, Cozzi A, Carla V, Sili M, Cavone L, et al. G-protein coupled receptor 35 (GPR35) activation and inflammatory pain: Studies on the antinociceptive effects of kynurenic acid and zaprinast. Neuropharmacology. Jun 2011;60(7-8):1227–31.

17. Zhang Y, Shi T, He Y. GPR35 regulates osteogenesis via the Wnt/GSK3beta/beta-catenin signaling pathway. Biochem Biophys Res Commun. Jun 4 2021;556:171–8.

18. Hansen MS, Madsen K, Price M, Soe K, Omata Y, Zaiss MM, et al. Transcriptional reprogramming during human osteoclast differentiation identifies regulators of osteoclast activity. Bone Res. Jan 24 2024;12(1):5.

19. Ma J, Chen P, Deng B, Wang R. Kynurenic acid promotes osteogenesis via the Wnt/β-catenin signaling. In Vitro Cellular and Developmental Biology - Animal. 2023;59(5):356–65.

20. Taniguchi Y, Tonai-Kachi H, Shinjo K. Zaprinast, a well-known cyclic guanosine monophosphate-specific phosphodiesterase inhibitor, is an agonist for GPR35. FEBS Lett. Sep 18 2006;580(21):5003–8.

21. Park SJ, Lee SJ, Nam SY, Im DS. GPR35 mediates lodoxamide-induced migration inhibitory response but not CXCL17-induced migration stimulatory response in THP-1 cells; is GPR35 a receptor for CXCL17? Br J Pharmacol. Jan 2018;175(1):154–61.

22. Jenkins L, Alvarez-Curto E, Campbell K, de Munnik S, Canals M, Schlyer S, et al. Agonist activation of the G protein-coupled receptor GPR35 involves transmembrane domain III and is transduced via Galpha(1)(3) and beta-arrestin-2. Br J Pharmacol. Feb 2011;162(3):733–48.

23. Schihada H, Klompstra TM, Humphrys LJ, Cervenka I, Dadvar S, Kolb P, et al. Isoforms of GPR35 have distinct extracellular N-termini that allosterically modify receptor-transducer coupling and mediate intracellular pathway bias. J Biol Chem. Sep 2022;298(9):102328.

24. Duan J, Liu Q, Yuan Q, Ji Y, Zhu S, Tan Y, et al. Insights into divalent cation regulation and G(13)-coupling of orphan receptor GPR35. Cell Discov. Dec 21 2022;8(1):135.

25. Hansen MS. Measuring Calcium Levels in Bone-Resorbing Osteoclasts and Bone-Forming Osteoblasts. Methods Mol Biol. 2025;2861:167–86.

26. Simonsen JL, Rosada C, Serakinci N, Justesen J, Stenderup K, Rattan SI, et al. Telomerase expression extends the proliferative life-span and maintains the osteogenic potential of human bone marrow stromal cells. Nat Biotechnol. Jun 2002;20(6):592–6.

27. Hansen MS, Soe K, Christensen LL, Fernandez-Guerra P, Hansen NW, Wyatt RA, et al. GIP reduces osteoclast activity and improves osteoblast survival in primary human bone cells. Eur J Endocrinol. Jan 10 2023;188(1).

28. Pfaffl MW. A new mathematical model for relative quantification in real-time RT-PCR. Nucleic Acids Res. May 1 2001;29(9):e45.

29. Jamaluddin A. Quantifying Gq Signaling Using the IP(1) Homogenous Time-Resolved Fluorescence (HTRF) Assay. Methods Mol Biol. 2025;2861:23–32.

30. Destaing O, Sanjay A, Itzstein C, Horne WC, Toomre D, De Camilli P, et al. The tyrosine kinase activity of c-Src regulates actin dynamics and organization of podosomes in osteoclasts. Mol Biol Cell. Jan 2008;19(1):394–404.

31. Moon JB, Kim JH, Kim K, Youn BU, Ko A, Lee SY, et al. Akt induces osteoclast differentiation through regulating the GSK3beta/NFATc1 signaling cascade. J Immunol. Jan 1 2012;188(1):163–9.

32. Li X, Udagawa N, Takami M, Sato N, Kobayashi Y, Takahashi N. p38 Mitogen-activated protein kinase is crucially involved in osteoclast differentiation but not in cytokine production, phagocytosis, or dendritic cell differentiation of bone marrow macrophages. Endocrinology. Nov 2003;144(11):4999–5005.

33. Lin J, Lee D, Choi Y, Lee SY. The scaffold protein RACK1 mediates the RANKL-dependent activation of p38 MAPK in osteoclast precursors. Sci Signal. Jun 2 2015;8(379):ra54.

34. Miyazaki T, Sanjay A, Neff L, Tanaka S, Horne WC, Baron R. Src kinase activity is essential for osteoclast function. J Biol Chem. Apr 23 2004;279(17):17660–6.

35. Iotsova V, Caamano J, Loy J, Yang Y, Lewin A, Bravo R. Osteopetrosis in mice lacking NF-kappaB1 and NF-kappaB2. Nat Med. Nov 1997;3(11):1285–9.

36. Gingery A, Bradley E, Shaw A, Oursler MJ. Phosphatidylinositol 3-kinase coordinately activates the MEK/ERK and AKT/NFkappaB pathways to maintain osteoclast survival. J Cell Biochem. May 1 2003;89(1):165–79.

37. Wong BR, Besser D, Kim N, Arron JR, Vologodskaia M, Hanafusa H, et al. TRANCE, a TNF family member, activates Akt/PKB through a signaling complex involving TRAF6 and c-Src. Mol Cell. Dec 1999;4(6):1041–9.

38. Du K, Montminy M. CREB is a regulatory target for the protein kinase Akt/PKB. J Biol Chem. Dec 4 1998;273(49):32377–9.

39. Oka S, Ota R, Shima M, Yamashita A, Sugiura T. GPR35 is a novel lysophosphatidic acid receptor. Biochem Biophys Res Commun. Apr 30 2010;395(2):232–7.

40. Lin LC, Quon T, Engberg S, Mackenzie AE, Tobin AB, Milligan G. G Protein-Coupled Receptor GPR35 Suppresses Lipid Accumulation in Hepatocytes. ACS Pharmacol Transl Sci. Dec 10 2021;4(6):1835–48.

41. Shi T, Shi Y, Gao H, Ma Y, Wang Q, Shen S, et al. Exercised accelerated the production of muscle-derived kynurenic acid in skeletal muscle and alleviated the postmenopausal osteoporosis through the Gpr35/NFkappaB p65 pathway. J Orthop Translat. Jul 2022;35:1–12.

42. Sundaram K, Nishimura R, Senn J, Youssef RF, London SD, Reddy SV. RANK ligand signaling modulates the matrix metalloproteinase-9 gene expression during osteoclast differentiation. Exp Cell Res. Jan 1 2007;313(1):168–78.

43. Quon T, Lin LC, Ganguly A, Hudson BD, Tobin AB, Milligan G. Biased constitutive signaling of the G protein-coupled receptor GPR35 suppresses gut barrier permeability. J Biol Chem. Jan 2025;301(1):108035.

44. Wyant GA, Yu W, Doulamis IP, Nomoto RS, Saeed MY, Duignan T, et al. Mitochondrial remodeling and ischemic protection by G protein-coupled receptor 35 agonists. Science. Aug 5 2022;377(6606):621-9.

