## Supplemental text for "G protein-coupled receptor 35 (GPR35) stimulation reduces osteoclast activity in primary human bone cells"

### Supplementary Appendix

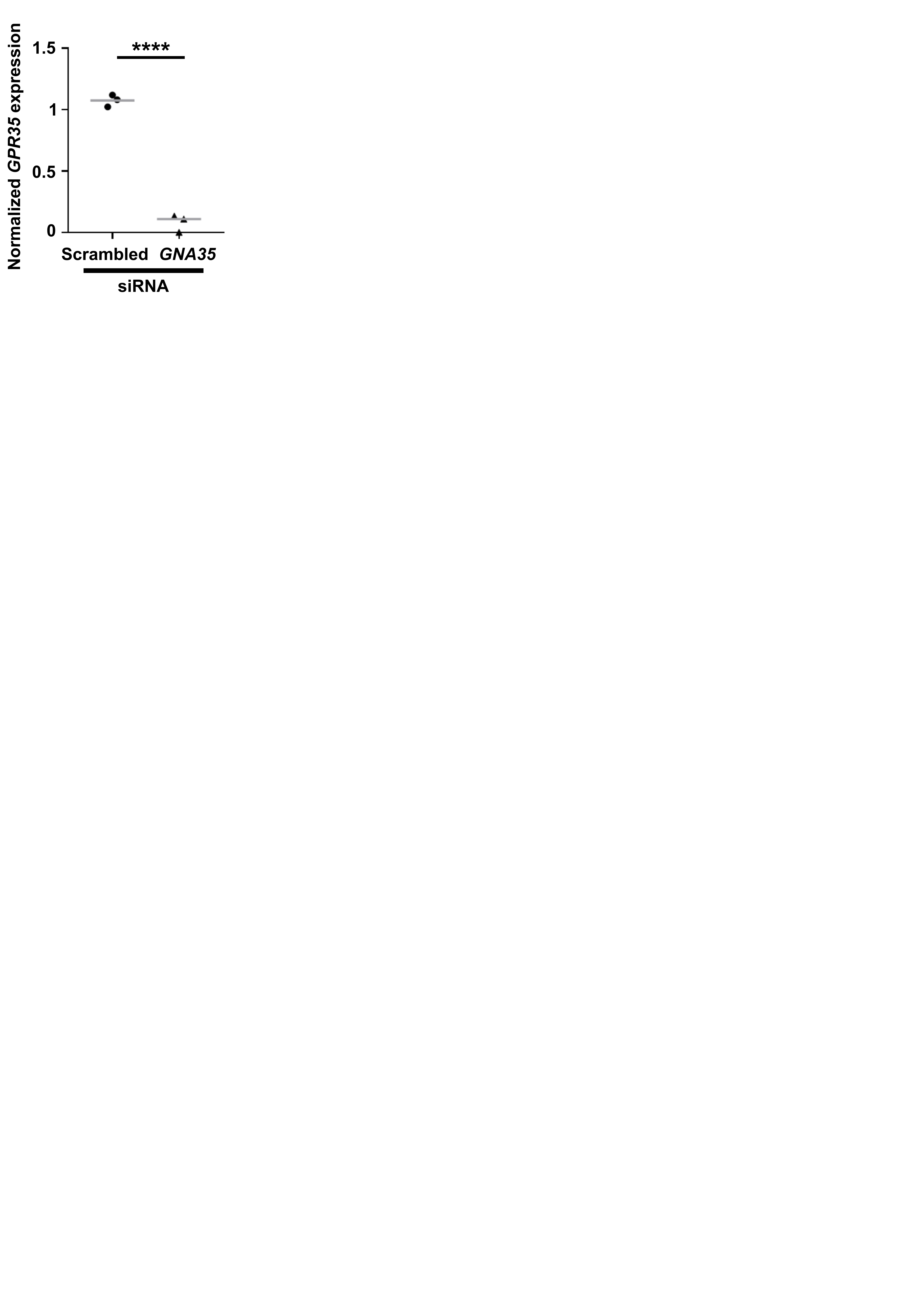

**Supplementary Figure 1	Validation that siRNA significantly reduces *GPR35* gene expression**

Gene expression of *GPR35* in mature osteoclasts exposed to scrambled or *GPR35-*targeted siRNAs determined by qPCR. Data was normalized to the geometric mean of three housekeeper genes (*ACTB, RPLP0, UBC*). Each point represents an independent donor. The gray line shows mean. Statistical analyses were performed by unpaired t-test. ****p<0.0001.

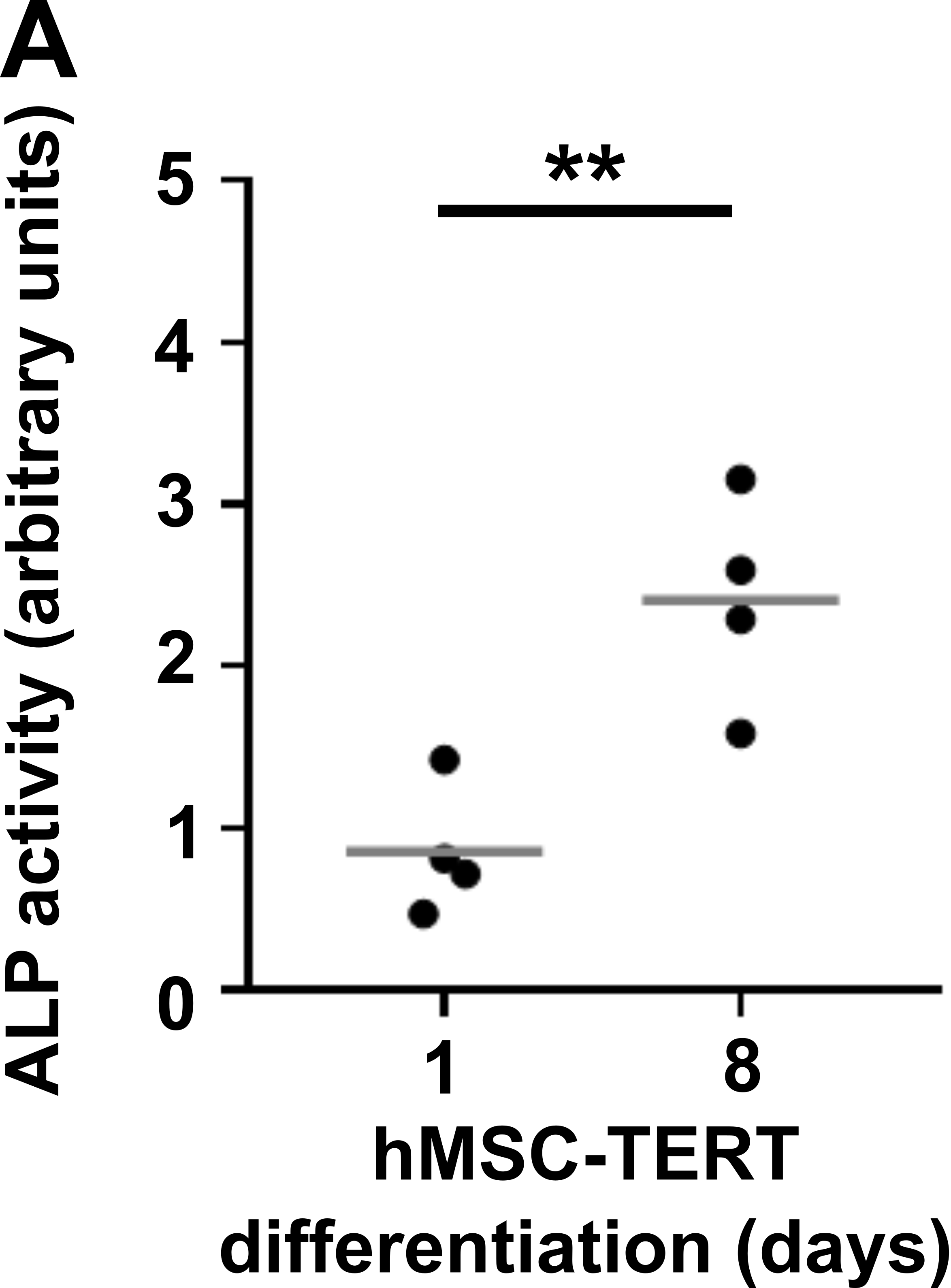

**Supplementary Figure 2	hMSC-TERT cells have increased alkaline phosphatase activity following differentiation**

Quantification of alkaline phosphatase activity in undifferentiated hMSC-TERT cells (day 1) and after eight days differentiation to osteoblasts. Each point represents an independent passage of cells. The gray line shows the mean. Statistical analysis was performed by unpaired t-test. **p<0.01.

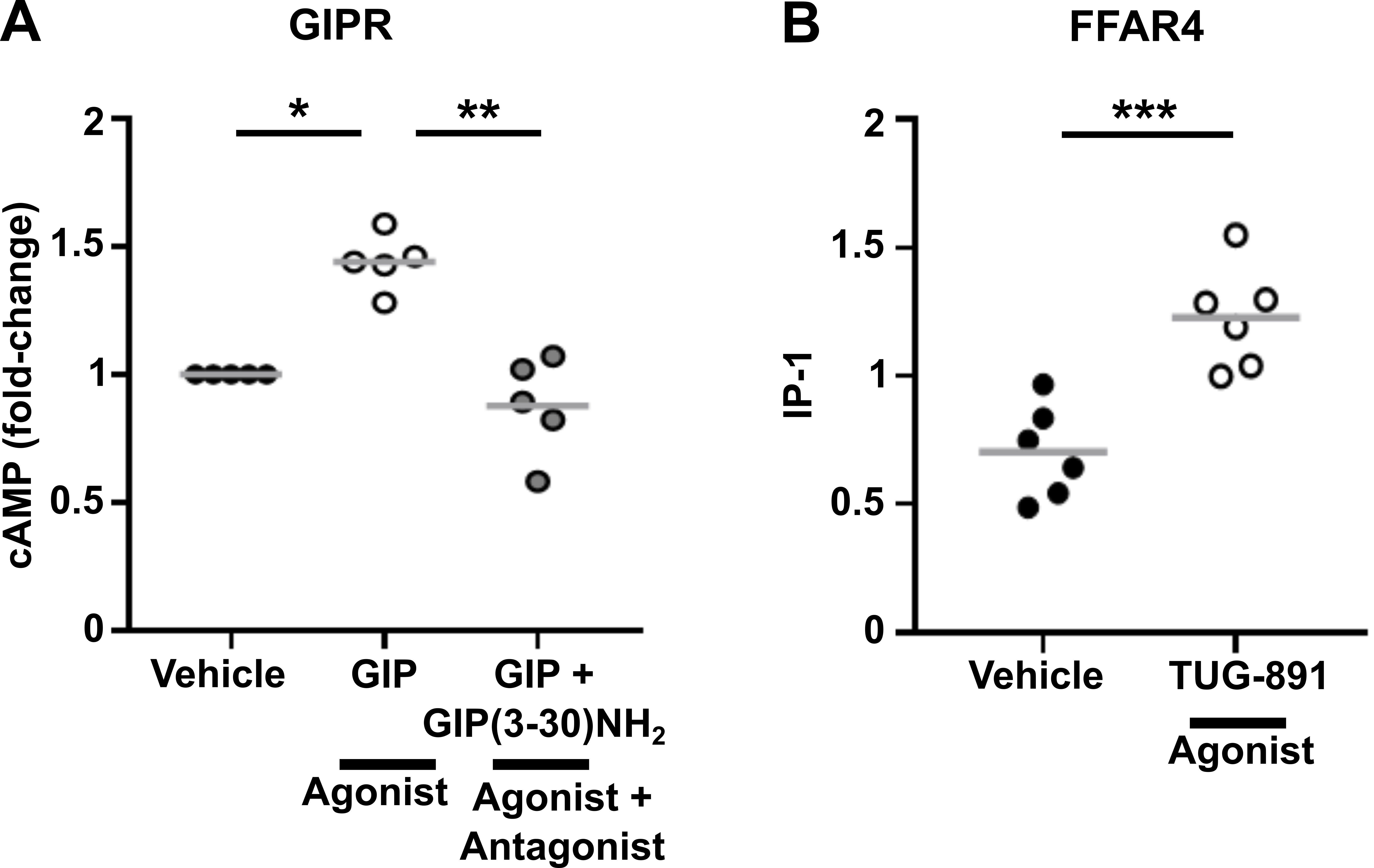

**Supplementary Figure 3	Confirmation that assays accurately measure Gs and Gq/11 activation**

(**A**) cAMP concentrations measured by LANCE cAMP assays in primary human osteoclasts exposed to vehicle, GIP or GIP with GIPR antagonist, GIP(3-30)NH_2_. GIPR is known to activate Gs-cAMP signaling^1^. (**B**) IP-1 concentrations measured by IP-one assays in primary human osteoclasts exposed to vehicle or TUG-891, an agonist for FFAR4 which we have previously shown activates Gq signaling^2^. The gray line shows median in A and mean in B. Each point represents an independent donor. Statistical analysis by Kruskal-Wallis test with Dunn’s multiple comparisons testing in A and unpaired t-test for B. ***p<0.001, ** p<0.01, *p<0.05.

#### Supplementary Table 1 Primers used in qPCR experiments

| **Gene Function** | **Gene name** | **Protein name** | **Gene Globe ID** |
| --- | --- | --- | --- |
| Housekeeper | *ACTB* | β-actin | QT00095431 |
|  | *RPLP0:* | Ribosomal protein lateral stalk subunit P0 | QT00075012 |
|  | *UBC* | Ubiquitin C | QT00234430 |
| Osteoclast activity | *CTSK* | Cathepsin K | QT00093856 |
|  | *MMP9* | Matrix metalloproteinase 9 | QT00040040 |
| G protein activity | *GNA11* | Gα11 | QT00084987 |
|  | *GNA12* | Gα12 | QT00235858 |
|  | *GNA13* | Gα13 | QT00079968 |
|  | *GNAQ* | Gαq | QT00037296 |
| GPCR | *GPR35* | G protein-coupled receptor 35 | QT02403128 |

**References**

1	Hansen, M. S. *et al.* GIP reduces osteoclast activity and improves osteoblast survival in primary human bone cells. *Eur J Endocrinol* **188** (2023). <https://doi.org:10.1093/ejendo/lvac004>

2	Hansen, M. S. *et al.* Transcriptional reprogramming during human osteoclast differentiation identifies regulators of osteoclast activity. *Bone Res* **12**, 5 (2024). <https://doi.org:10.1038/s41413-023-00312-6>
